# Fishing for New Bt Receptors in Diamondback Moth

**DOI:** 10.1101/181834

**Authors:** Yazhou Chen, Yuping Huang, Qun Liu, Jun Xu, Saskia Hogenhout, Yongping Huang, Anjiang Tan

**Author notes:** Correspondence should be addressed to YH and AT.

## Abstract

Bt toxins bind to receptors in the brush border membrane of the insect gut and create pores, leading to insect death. Bt-resistant insects demonstrate reduced binding of the Bt toxins to gut membranes. However, our understanding of the gut receptors involved in Bt toxin binding, and which receptors confer resistance to these toxins is incomplete, especially in diamondback moth (*Plutella xylostella*), a major agricultural pest. Identifying receptors has remained challenging because we lack sufficiently sensitive methods to detect Bt receptor interactions. Here, we report a modified far-immunoblotting technique, which revealed a broad spectrum of binding targets for the Bt toxins Cry1Ac, Cry1Ab, and Cry1Bd in diamondback moth. We confirm the role of the glucosinolate sulfatases GSS1 and GSS2 in Cry1Bd toxicity. GSS1 and GSS2 bind directly to Cry1Bd, and their expression is crucial for Cry1Bd toxicity. These results improve our understanding of the molecular mechanisms of Bt toxicity.

**AUTHOR SUMMARY:** The Bt toxins, from the soil bacterium *Bacillus thuringiensis*, have wide applications in agriculture as insecticides applied to plants or expressed in genetically modified crops. Bt toxins bind to receptors in the brush border membrane of the insect gut and create pores leading to insect death. The success of the Bt toxins in controlling insect pests has been hindered by the emergence of resistant insects, which show reduced binding of Bt to their gut membranes. Although ongoing research has identified a few receptors, many remain unknown and the mechanisms by which these receptors cause resistance remain unclear. Here, we used a modified far-immunoblotting technique to identify proteins that bind to the toxins Cry1Ac, Cry1Ab, and Cry1Bd in the diamondback moth. This identified two glucosinolate sulfatases that bind directly to Cry1Bd; also, the toxicity of Cry1Bd requires expression of these glucosinolate sulfatases. Therefore, identification of these candidate receptors improves our understanding of Bt function and resistance.

## INTRODUCTION

*Bacillus thuringiensis* (Bt) is a Gram-positive, soil-dwelling bacterium that produces δ-endotoxin proteins known as Bt toxins or Cry toxins (crystalline toxins). Bt toxins efficiently kill lepidopteran, dipteran, and coleopteran pests [1], but are harmless to humans and other vertebrate animals [1]. The Bt toxins belong to a class of bacterial pore-forming toxins. Once ingested by insects, Bt protoxins are solubilized in the insect midgut, and are then cleaved by proteases to produce activated toxins [2]. These activated toxins penetrate the insect midgut protrophic membrane and bind to specific target sites, called primary receptors (such as cadherin), of the brush border membrane vesicles (BBMV) [1, 3]. Interactions between Bt toxins and cadherin facilitate protease cleavage of the helix α-1 of the toxin, promoting toxin oligomerization [4]. These toxin oligomers are thought to have increased binding affinity to secondary receptors including glycosylphosphatidylinositol (GPI)-anchored proteins, aminopeptidase N (APN) [5], and alkaline phosphatases (ALP) [6]. Binding of the toxin oligomers to these secondary receptors creates pores in the midgut membranes, thus causing osmotic shock, breakdown of the midgut cells, and insect death [4–6]. However, others have proposed that binding of the activated Bt toxin monomers to cadherin initiates a magnesium-dependent signaling pathway, causing cell disruption [7]. In either model, the binding of Bt toxins to various midgut receptors is essential for disrupting the midgut membrane, which leads to cell lysis.

Insects develop resistance to Bt toxins by evolving mechanisms that reduce or interfere with the ability of the Bt toxins to bind to receptors [8]. To date, seven insect species commonly found in open field and greenhouse crops have developed resistance to Cry toxins [9]. It is important to understand the molecular mechanisms of toxin action, and identify the genes contributing to insect resistance, to develop strategies for the long-term and sustainable use of Bt and their Cry toxins as insecticides.

The diamondback moth (DBM) *Plutella xylostella* (Lepidoptera: Plutellidae) causes US $4–5 billion in annual management costs [10] and is the first insect that was reported to have evolved resistance to Bt toxins in open fields [11]. The DBM resistance phenotype involves reduced binding of toxins to the brush border membrane cell proteins, a trait that is inherited in a recessive manner but which achieves high resistance levels [12, 13]. Multiple Bt toxin receptors have been identified in lepidopteran insects [14]. However, genetic analysis has conclusively eliminated these as conferring resistance to Cry1A in DBM. Resistance mechanisms may involve alterations in the expression levels of Bt toxin receptors [15, 16]. For example, down-regulation of ALP and adenosine triphosphate (ATP)-binding cassette transporter subfamily C (ABCC) gene expression has been linked to DBM resistance to the Bt toxin Cry1Ac [16]. Comparing the sequences of these genes between susceptible and resistant DBM strains revealed no obvious mutations to explain the Cry1Ac resistance phenotype, thus casting doubt on the role of these proteins [16].

Finding Bt toxin receptors in insect midguts has been challenging, partly because there are no sufficiently sensitive methods to detect Bt receptor interactions. Whereas genetics-based methods have identified cadherin and ABCC2 genes as being associated with Bt resistance in *Heliothis virescens* and *Bombxy mori* [17–19], their involvement in DBM resistance to Bt toxins is unclear [20, 21]. Current methods are unlikely to identify low abundance membrane proteins, which could contribute to Bt toxin function [22, 23].

Here, we report the use of a modified far-immunoblotting method to identify 81 candidate insect proteins that interact with the Bt toxins Cry1Ac, Cry1Ab, and Cry1Bd in the DBM BBMV. In addition to cadherin and APN2, we identified two glucosinolate sulfatase proteins (GSS1 and GSS2) that interact with Cry1Bd. These GSSs were previously shown to protect DBM against glucosinolates from Brassicaceae and likely degrade these toxic plant compounds [24]. Follow-up work confirmed a crucial role for GSSs in Bt toxin activity. Our study provides a novel method to identify insect proteins that interact with Bt toxins. Moreover, we discovered new components that contribute to the action of Bt toxins in the insect midgut. Taken together, this work advances our ability to uncover mechanisms involved in Bt toxin action and resistance.

## RESULTS

### Binding spectrum of Bt toxins

We first confirmed the susceptibility of the DBM strain ‘Fuzhou’ to three Lepidoptera-specific Bt toxins, Cry1Ac, Cry1Ab, and Cry1Bd by determining the concentration of the toxin that was lethal to 50% of the DBM (LC50). The LC50s of Cry1Ac, Cry1Ab, and Cry1Bd against the third larvae of this strain were 4.35 mg/L, 0.49 mg/L, and 0.05 mg/L, respectively, indicating that the Fuzhou strain is highly susceptible to Cry1Ac, Cry1Ab, and Cry1Bd, as reported previously [13, 25–29].

It has been proposed that Bt toxin receptors are located on the midgut epithelium, where activated toxins bind with receptors and form pore structures that insert into the membrane [30]. To detect unknown receptors contributing to DBM susceptibility to Bt toxins, we developed a modified far-immunoblotting method to detect interactors based on the presence or absence of Bt toxin binding sites. Each Bt toxin (Cry1Ac, Cry1Ab, and Cry1Bd) was separated by sodium dodecyl sulfate polyacrylamide gel electrophoresis (SDS-PAGE), and transferred onto nitrocellulose membrane (Supplemental Figure 1). Nitrocellulose membranes with the Bt toxin bands were cut, denatured and renatured by gradually reducing the guanidine-HCl concentration [31]. The membrane was then blocked with protein-free buffer and incubated with total DBM BBMV proteins to capture Cry toxin-binding insect proteins [31]. Nitrocellulose membrane sections containing Bt toxin–protein complexes were subjected to trypsin digestion. Digested peptides were dried in a vacuum, and the membrane was removed by adding acetone [32]. Precipitated peptides were air-dried and determined by nano liquid chromatography–tandem mass spectroscopy (nano LC-MS/MS) coupled with the Q-Exactive Orbitrap mass spectrometer (ThermoFisher).

**Fig 1.**
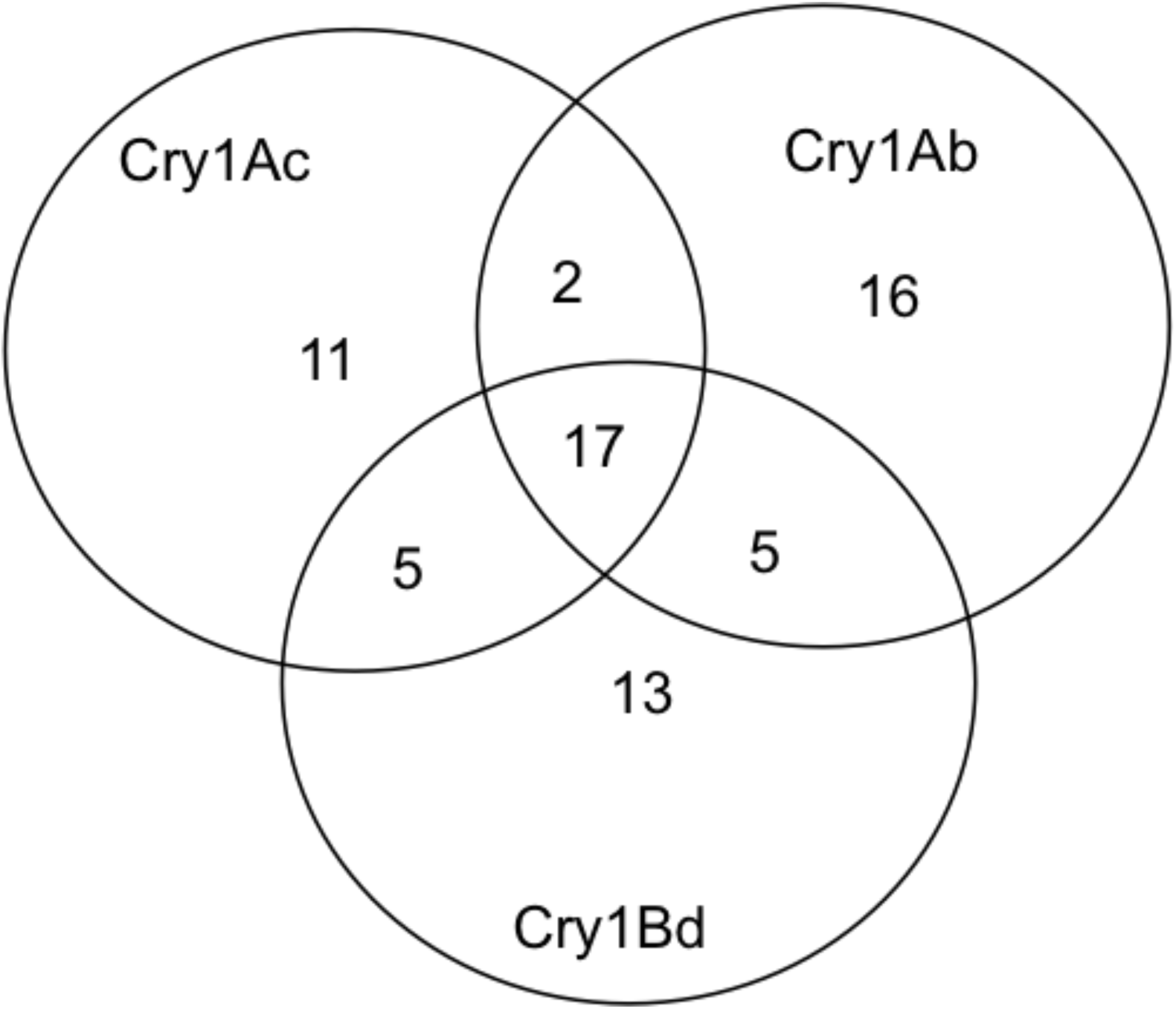
Venn diagram showing proteins identified by the modified far-immunoblotting method.

**Supplemental Fig 1.** A flowchart to describe the modified far-immunoblotting method used in this study.

MS/MS spectra were queried against a combined protein database including DBM protein sequences and protein sequences of Cry1Ac, Cry1Ab, and Cry1Bd. A total of 520 peptides were detected in three bands. In each band, the Bt toxin was the primary protein based on number of peptides identified (Table 1). In the Cry1Ac sample, 38.2% of peptides were identified as Cry1Ac, 26.5% as Cry1Ab in Cry1Ab sample, and 27.6% as Cry1Bd in Cry1Bd sample (Supplemental Table 1). The rest of the peptides corresponded to 81 unique proteins with at least one peptide, and had a wide range of masses and isoelectric points (pIs) (Supplemental Figure 2). This implies that the modified far-immunoblotting method captured targets from the proteome scale. Of these, 35 were in the Cry1Ac band, 40 were in the Cry1Ab band, and 39 were in the Cry1Bd band (Table 1, Figure 1). Besides Bt toxins, the most abundant proteins captured from all samples were acetylcholinesterase, actin, adenosylhomocysteinase, and ryanodine receptor 44F (Supplemental Table 1). Cadherin and APN2, previously shown to interact with CryA [20, 33, 34], were also identified from three samples, thus providing support for the pore-formation model [1]. Some proteins also bound specifically to the Cry1Ac and Cry1Ab toxins, and each toxin captured several proteins in particular (Figure 1, Supplemental Table 1). Future research might investigate whether loss of binding to those proteins contributes to the development of DBM resistance to Cry1Ac and Cry1Ab toxins.

**Table 1.** Peptide identification analysis on a Q-Exactive Orbitrap LC-MS system (60 min gradient)

| Bt Protein | MS Spectra | MSMS Spectra | Identification (%) | Identified Peptides |  |  | Unique Peptides |  | Proteins | Isotope pattern |
| --- | --- | --- | --- | --- | --- | --- | --- | --- | --- | --- |
|  |  |  |  | Total | Bt | % | Total | Bt |  |  |
| Cry1Ac | 5946 | 16282 | 0.83 | 136 | 52 | 38.2% | 60 | 17 | 35 | 34871 |
| Cry1Ab | 5837 | 17088 | 1.29 | 189 | 49 | 25.9% | 95 | 20 | 40 | 36438 |
| Cry1Bd | 5820 | 16797 | 0.97 | 163 | 45 | 27.6% | 64 | 11 | 39 | 33664 |

| Bt<br>Protein | MS<br>Spectra | MSMS<br>Spectra | Identification<br>(%) | Identified Peptides |  |  | Unique Peptides |  | Proteins | Isotope pattern |
| --- | --- | --- | --- | --- | --- | --- | --- | --- | --- | --- |
|  |  |  |  | Total | Bt | % | Total | Bt |  |  |
| Cry1Ac | 5946 | 16282 | 0.83 | 136 | 52 | 38.2% | 60 | 17 | 35 | 34871 |
| Cry1Ab | 5837 | 17088 | 1.29 | 189 | 49 | 25.9% | 95 | 20 | 40 | 36438 |
| Cry1Bd | 5820 | 16797 | 0.97 | 163 | 45 | 27.6% | 64 | 11 | 39 | 33664 |

**Fig 2.**
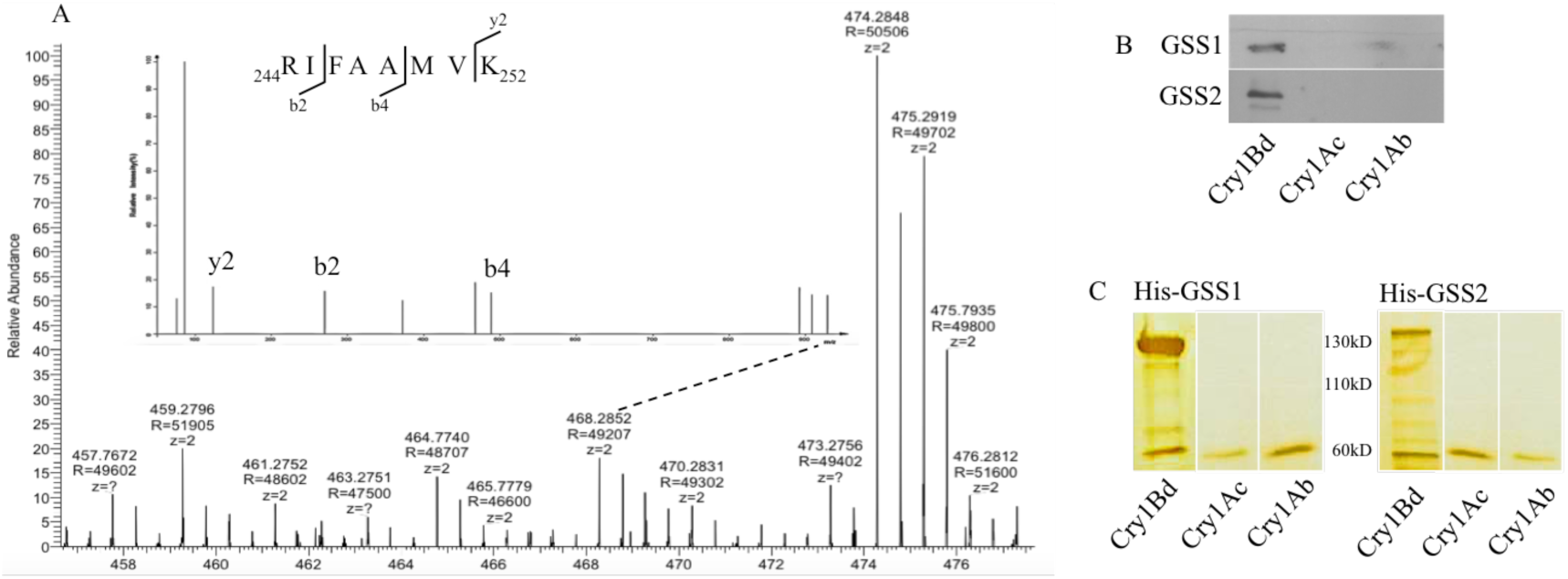
Binding of GSSs to Cry1Bd. A. The peptide ^244^ RIFAAMVK ^252^, identified by MS in the Cry1Bd sample, matched GSS1 and GSS2. B. Binding of GSS1 and GSS2 to Cry1Bd was detected by far-immunoblotting. Bt proteins on nitrocellulose membranes were denatured and renatured by gradually reducing the guanidine-HCl concentration, then incubated with 5 μg His-GSS1 or His-GSS2 after the membrane was blocked. Anti-His antibody was used to detect binding by recognizing His-tags fused with GSS1 or GSS2 to Cry1Bd but not Cry1A or Cry1Ab. C. Binding of GSS1 and GSS2 to Cry1Bd was detected by pull-down assay. His-GSS1 or His-GSS2 bound to cobalt resin was incubated with 150 μg of one Bt protein. Cry1Ac or Cry1Ab was removed using washing buffer, but Cry1Bd remained and was co-eluted with GSS1 or GSS2 by elution buffer.

**Table 1. Peptide identification analysis on a Q-Exactive Orbitrap LC-MS system**

**Fig 1. Venn diagram showing proteins identified by the modified far-immunoblotting method**

**Supplemental Table 1.** List of proteins identified from Cry1Ac, Cy1Ab, and Cry1Bd samples.

**Supplemental Fig 2.** Molecular range (A) and pH (B) of proteins detected from Cr1Ac, Cry1Ab, and Cry1Bd using the modified far-immunoblotting method.

Closer attention was paid to the proteins captured by Cry1Bd. This Cry toxin has the greatest potential to be used against DBM because the field population of this insect has evolved cross-resistance to four Bt toxins including Cry1Ac and Cry1Ab, but remains highly susceptible to Cry1Bd [13, 25, 27–29, 35]. Therefore, if Cry1Bd is to be used to control this pest, it is important to investigate potential Cry1Bd receptors [36, 37].

### Glucosinolate sulfatases are Cry1Bd binding sites

Since Cry1Bd has unique binding sites that do not interact with Cry1Ac or Cry1Ab [8, 25, 36, 38], we focused on proteins captured only by Cry1Bd. Bt toxin receptors are characterized as transmembrane proteins, like the primary receptor cadherin, or as GPI-anchored proteins such as the secondary receptors ALP and APN. We analyzed the transmembrane helices using the Transmembrane Hidden Markov Model server (TMHMM) [39] and the GPI-anchor sequences of the candidates using GPI Modification Site Prediction [40]. This revealed four candidates with GPI-anchor sites or transmembrane helixes: ATP synthase F0 subunit 8, β-1,3-glycosyltransferase 5, and two glucosinolate sulfatases. The mitochondrial protein ATP synthase F0 subunit 8 has been eliminated as a Bt toxin target [41, 42]; likewise, β-1,3-glycosyltransferase 5 has been ruled out as contributing to Bt resistance in *Plutella* [43]. Two glucosinolate sulfatases (GSS), GSS1 and GSS2, matched one peptide ^244^RIFAAMVK^252^ (Figure 2A). GSS1 and GSS2 share 96% amino acid identity and lie adjacent to each other in the DBM genome (Supplemental Figure 3). Both have N-terminal secretory signal peptides, indicating that they are secreted in insect midgut cells (Supplemental Figure 4). In addition, GSS2 contains an N terminal transmembrane helix, indicating that it is a membrane anchor protein with a C-terminal extending outside of the cell membrane (Supplemental Figure 4). Both GSSs have predicted GPI-anchor sites at C485 (Supplemental Figure 5), indicating their potential as Cry1Bd binding sites.

**Fig 3.**
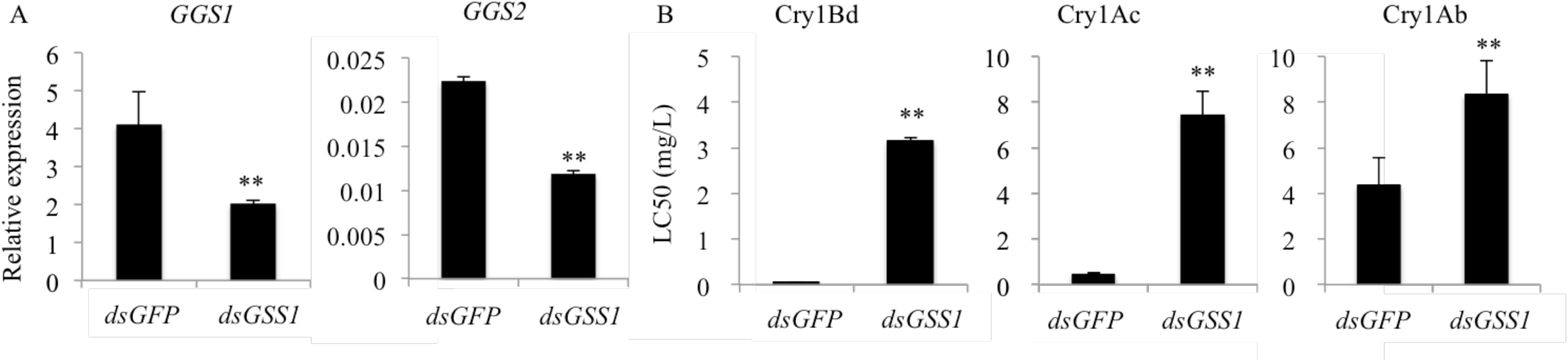
Reduction of expression of *GSS1* and *GSS2* results in increased tolerance of DBM to Bt toxins. A. Expression levels of *GSS1* and *GSS2* in *GSS*-silenced larvae. Freshly hatched DBM larvae were fed leaves of either *dsGFP* or *dsGSS1* lines for about 7 d until larvae reached the third instar, whereupon they were harvested for RNA analysis. Values (mean ± SD) were obtained from three independent experiments. ** above the columns indicates statistical significance between samples (*P*<0.01). B. LC50 of *GSS*-silenced larvae against Bt toxins. Two leaves taken from 4-week-old *A. thaliana* plants of genotypes *dsGFP* or *dsGSS1* were laid on a moistened filter paper in a 15 mm petri dish. Freshly hatched DBM larvae were placed on leaves of each genotype and fed for 7 d. On the eighth day, larvae were transferred to fresh leaves coated with a diluted suspension of Bt proteins in HEPES buffer (pH 8.0), or HEPES buffer as a control. Mortality was recorded after 24 h and LC50 was calculated by probit analysis based on the dose determined to be high enough to kill 100% of larvae. Values (mean ± SD) were obtained from three independent experiments. ** above the columns indicates statistical significance between samples (*P*<0.01).

**Fig 4.**
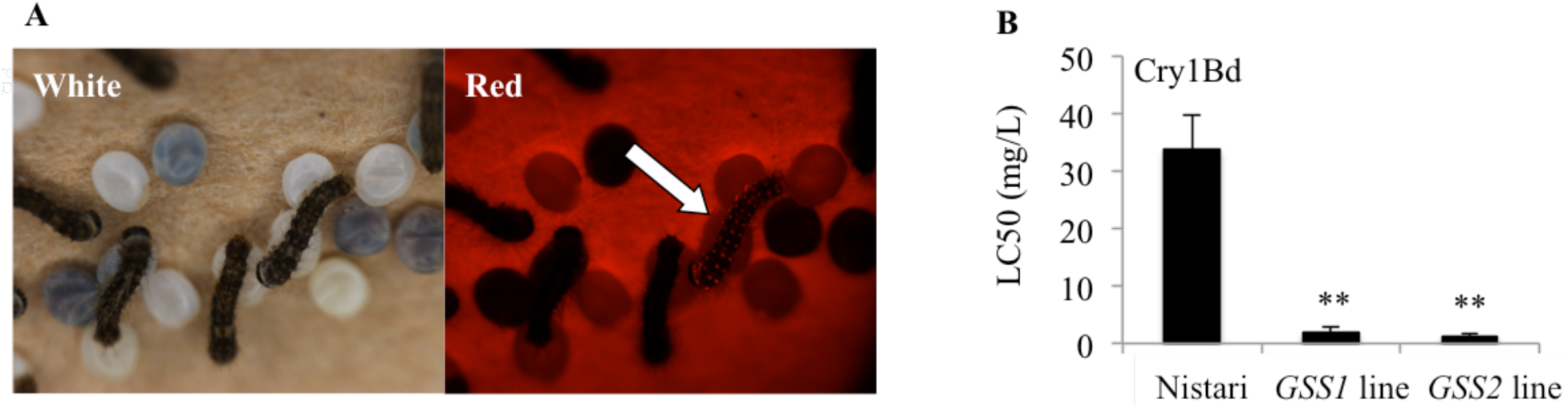
Transgenic silkworms expressing DBM *GSS1* or *GSS2* became susceptible to Cry1Bd. A. DsRed fluorescence phenotypes of hybrid offspring produced by transformed silkworms and wild type Nistari. Insects with DsRed fluorescence (indicated by arrow) were selected for Cry1Bd toxicity tests. B. LC50 of hybrid silkworm against Cry1Bd. Four squares of mulberry leaf (4 × 4 cm), coated with 50 μL of a diluted suspension of Cry1Ab protoxin in HEPES buffer (pH 8.0), were fed to 10 second instar larvae for 24 h. The dose high enough to kill 100% of susceptible larvae was determined. Probit analysis was carried out using SPSS to determine the LC50 value. Values (mean ± SD) were obtained from three independent experiments. ** indicates statistical significance between samples (*P*<0.01).

**Fig 2. Binding of GSSs to Cry1Bd**

A. The peptide ^244^RIFAAMVK^252^, identified by MS in the Cry1Bd sample, matched GSS1 and GSS2.

B. Binding of GSS1 and GSS2 to Cry1Bd was detected by far-immunoblotting. Bt proteins on nitrocellulose membranes were denatured and renatured by gradually reducing the guanidine-HCl concentration, then incubated with 5 μg His-GSS1 or His-GSS2 after the membrane was blocked. Anti-His antibody was used to detect binding by recognizing His-tags fused with GSS1 or GSS2 to Cry1Bd but not Cry1A or Cry1Ab.

C. Binding of GSS1 and GSS2 to Cry1Bd was detected by pull-down assay. His-GSS1 or His-GSS2 bound to cobalt resin was incubated with 150 μg of one Bt protein. Cry1Ac or Cry1Ab was removed using washing buffer, but Cry1Bd remained and was co-eluted with GSS1 or GSS2 by elution buffer.

**Supplemental Fig 3.** Structure of *GSS1* and *GSS2* genes

The *GSS1* locus is next to a highly similar paralog *GSS2*. Arrows indicate direction of transcription; boxes in grey indicate exons.

**Supplemental Fig 4.** Transmembrane helices of GSS1 (top) and GSS2 (bottom) analyzed by TMHMM.

Both GSS1 and GSS2 have N-terminal secretory signal peptides, indicating that they are extracellular proteins. GSS2 also contains a transmembrane helix at its N-terminal, indicating that it is a membrane anchor protein with a C-terminal extending outside of the cell membrane.

**Supplemental Fig 5.** GPI-anchor sites of GSS1 (A) and GSS2 (B) predicted by GPI Modification Site Prediction [40].

In the previous experiment, Bt toxins were incubated with whole BBMV preparations, which may contain protein complexes that are able to bind the Bt toxins. To test whether GSSs bind Bt toxins directly, or as part of a larger protein complex, we incubated the Bt toxins with His-tagged GSS1 and GSS2 alone (using the far-immunoblotting technique; see Materials and Methods). This showed that both GSSs bind Cry1Bd but not Cry1Ab or Cry1Ac; His-tag pull-down experiments further confirmed these results (Figure 2C), indicating that the interaction between GSSs and Cry1Bd is direct and specific.

### GSSs are critical for Cry1Bd toxicity

GSSs are enzymes used by DBM to protect itself against the accumulation of toxic compounds from Brassicaceae [24]. When these plants are damaged by herbivory, a myrosinase processes glucosinolates into compounds that are toxic to the insect [44]. DBM counters this process by using GSSs to convert glucosinolates into non-toxic compounds and sulfate, which inhibits myrosinase activity in the plant [24]. We hypothesized that GSSs are targets of Cry1Bd and critical for Cry1Bd toxicity to DBM. To investigate this possibility, we generated transgenic Arabidopsis plants expressing *dsGSS1*, which silences both *GSS* genes because of their highly similar sequences. Freshly hatched DBM larvae were fed leaves of *dsGFP* or *dsGSS1* lines for about 7 d until the third instar, when they were harvested for RNA analysis. *GSS*-silenced larvae showed no significant defects in body or fecal weight, which might be caused by functional compensation by other sulfatases (Supplemental Table 2). As shown in Figure 3A, expression levels of target genes decreased in those larvae. Third-instar DBM larvae were fed *dsGSS1* leaves coated with Bt toxin for 24 h, or with *dsGFP* leaves coated with the same Bt toxin as controls. *GSS*-silenced larvae had LC50 of 3.162 mg/L against Cry1Bd, an approximately 69-fold increase compared with controls (Figure 3B). *GSS*-silenced larvae also had a 15-fold increase in LC50 against Cry1Ab, and a 1.9-fold increase in LC50 against Cry1Ac (Figure 3B). These results show that GSSs are critical for Cry1Bd toxicity to DBM.

**Fig 3. Reduction of expression of *GSS1* and *GSS2* results in increased tolerance of DBM to Bt toxins**

A. Expression levels of *GSS1* and *GSS2* in *GSS*-silenced larvae. Freshly hatched DBM larvae were fed leaves of either *dsGFP* or *dsGSS1* lines for about 7 d until larvae reached the third instar, whereupon they were harvested for RNA analysis. Values (mean ± SD) were obtained from three independent experiments. ** above the columns indicates statistical significance between samples (*P*<0.01).

B. LC50 of *GSS*-silenced larvae against Bt toxins. Two leaves taken from 4-week-old *A. thaliana* plants of genotypes *dsGFP* or *dsGSS1* were laid on a moistened filter paper in a 15 mm petri dish. Freshly hatched DBM larvae were placed on leaves of each genotype and fed for 7 d. On the eighth day, larvae were transferred to fresh leaves coated with a diluted suspension of Bt proteins in HEPES buffer (pH 8.0), or HEPES buffer as a control. Mortality was recorded after 24 h and LC50 was calculated by probit analysis based on the dose determined to be high enough to kill 100% of larvae. Values (mean ± SD) were obtained from three independent experiments. ** above the columns indicates statistical significance between samples (*P*<0.01).

**Supplemental Table 2.** Sulfatases annotated using the Diamondback Moth Genome Database (DMB-DB) [52].

### GSSs are causative agents of Cry1Bd susceptibility

To confirm that DBM GSSs are causative agents of susceptibility to Cry1Bd, we introduced *GSS1* and *GSS2* into another lepidopteran insect, the silkworm *Bombyx mori*, because of a lack of genetic tools in DBM. The Bt-resistant strain Nistari [15] was selected, and two transgenic silkworm strains were established: one expressing *GSS1* and the other expressing *GSS2*, both expressing the fluorescent protein DsRed as a selectable marker. Inverse PCR of genomic DNA revealed that GSS1 strains had one copy of the transgene on chromosome 23, and GSS2 strains had one copy on chromosome 11 (Supplemental Figure 6). Since reduction of *GSS1* or *GSS2* leads to increased Cry1Bd resistance, we assumed that presence of a single allele of each gene would change the susceptibility of transgenic strains. Positive individuals were crossed with wild-type silkworms, and hybrid offspring possessing the target gene (identified by expression of DsRed at the larval stage) were selected (Figure 4A). All fluorescent hybrids expressed only one allele of the target gene.

**Fig 4. Transgenic silkworms expressing DBM *GSS1* or *GSS2* became susceptible to Cry1Bd**

A. DsRed fluorescence phenotypes of hybrid offspring produced by transformed silkworms and wild type Nistari. Insects with DsRed fluorescence (indicated by arrow) were selected for Cry1Bd toxicity tests.

B. LC50 of hybrid silkworm against Cry1Bd. Four squares of mulberry leaf (4 × 4 cm), coated with 50 μL of a diluted suspension of Cry1Ab protoxin in HEPES buffer (pH 8.0), were fed to 10 second instar larvae for 24 h. The dose high enough to kill 100% of susceptible larvae was determined. Probit analysis was carried out using SPSS to determine the LC50 value. Values (mean ± SD) were obtained from three independent experiments. ** indicates statistical significance between samples (*P*<0.01).

**Supplemental Fig 6.** Genomic insertion of *GSS1* (A) and *GSS2* (B).

Genomic insertion of *GSS1* and *GSS2* in transgenic silkworm lines, as revealed by inverse PCR and sequencing. The transgene integration site in *GSS1* lies in chromosome 23, between two genes KAIKOGA026485 and KAIKOGA026486. The transgene integration site in *GSS2* lies in chromosome 11, between BMgn01916 and BMgn011917. Chromosome localization and partial genomic DNA sequences between the Sau3AI site and the 3□ or the 5□ insert boundaries of the vector are shown. In all insertions, the TTAA insertion site found in canonical *piggyBac* insertions was found at the 3□ and 5□ insert boundary.

Resistance levels of the hybrids were tested at the second instar by feeding the larvae with Cry1Bd toxin-coated mulberry leaf discs and recording mortality after 24 h. Wild type Nistari had an LC50 of 33.90 mg/L; however, expression of DBM *GSS1* and *GSS2* resulted in a drop in LC50 to 1.86 mg/L and 1.30 mg/L, respectively (Figure 4B). These results show that presence of GSS1 and GSS2 increases the susceptibility of silkworm to Cry1Bd toxins.

## DISCUSSION

Disruption of receptor binding is the most common mechanism by which insects gain resistance to Bt toxins [8], but identifying the receptors that confer resistance is a challenge. Here we report a novel, modified far-immunoblotting method that allowed us to identify previously unknown Bt toxin receptors. This method will help researchers to identify Bt toxin receptors in insect-species for which we lack sufficient genomic information. With the help of mass spectrometry, identified proteins can be annotated using a protein database homolog search. This method will facilitate assessment of the risk of evolution of insect resistance to a particular Bt toxin under consideration for use in the field, and inform the choice of appropriate toxins to delay resistance.

In agreement with the pore-formation model [1], our modified far-immunoblotting technique revealed that the well-characterized receptors cadherin and APN2 are captured by Cry1Ac, Cry1Ab, and Cry1Bd. Furthermore, it revealed broader interactions between Bt toxins and their targets, which may be helpful in rethinking long-standing hypotheses and designing testable experiments. For example, the binding site modification hypothesis proposed four Bt toxin-binding sites to explain DBM cross-resistance to different Bt toxins [8]. Site 1 binds Cry1Aa only, and Site 2 binds Cry1Ab, Cry1Ac, and Cry1F [8]. Sites 3 (Cry1B-binding) and 4 (Cry1C-binding) are distinct and do not interact with other toxins. Resistance to a certain Bt toxin is dependent on modification of a site. For instance, modification of site 2 can abolish binding to Cry1Aa, Cry1Ab, Cry1Ac, and causes DBM cross-resistance to those Cry toxins [8]. In this study, arylphorin was detected by Cry1Ac and Cry1Ab in the susceptible DBM strain, but not by Cry1Bd, indicating that arylphorin may bind to site 2. Indeed, increased arylphorin expression in *Spodoptera exigua* correlated with *B. thuringiensis* resistance [45]. Arylphorin might be an important component of DBM resistance to multiple Cry toxins, including Cry1Ac and Cry1Ab. Future investigations into whether nonsynonymous substitutions in the arylphorin gene sequence cause DBM cross-resistance to multiple Bt toxins are warranted.

Our modified far-immunoblotting method suggested the existence of a set of proteins that specifically bind certain Bt toxins. We chose to further investigate those proteins captured by Cry1Bd because the DBM field population remains highly susceptible to Cry1Bd, yet shows cross-resistance to Cry1Ac and Cry1Ab. DBM GSSs have predicted N-terminal secretory signal peptides and GPI-anchored sites, implying that GSSs are extracellular proteins that are selectively included in lipid rafts when pore-forming toxins interact with their targets [2]. Binding experiments revealed that GSSs serve as Cry1Bd binding sites. Moreover, Arabidopsis-mediated RNAi analysis and *B. mori* transformation experiments confirmed the role of GSSs in Cry1Bd toxicity. GSSs have been found in higher levels in Cry1Ac-resistant strains [46]. However, binding of GSSs to Cry1Ac has been inconsistently reported, and this has led to different scenarios. For example, binding of GSSs to Cry1Ac was detected in resistant DBM, leading to a hypothesis that GSS sequesters Cry1Ac in resistant animals [47]. In contrast, binding of GSSs to Cry1Ac was not detected in a resistant strain (NO-QA) or a susceptible strain (Geneva 88) [46]. This yielded the hypothesis that, in Bt resistant DBM, GSS may be involved in stress responses [46]. In our binding experiments, GSSs bound specifically with Cry1Bd but not with Cry1Ac or Cry1Ab, supporting the second scenario. However, *GSS*-silenced larvae showed increased resistance against Cry1Ac and Cry1Ab (Figure 3B), implying that GSSs might be indirectly involved in the action of Cry1A toxins. In eukaryotes, sulfatases are extensively glycosylated before being transported to their destinations [48]. It is possible that GSSs might be involved in Cry1A toxicity through their terminal GalNAc residue. Indeed, GalNAc has been shown to bind with the carbohydrate-binding sites of domain III of Cry1Ac [49]. An alternative explanation is that GSSs mediate Cry1A toxicity via a signaling pathway. Sulfatases have been attributed pivotal roles in Wnt [50] and pheromone signaling [51]. Recently studies have revealed that the MAPK signaling pathway manipulates the expression of multiple receptors relating DBM resistance to Cry1Ac [16]. It will be important for future studies to investigate whether MAPK signaling pathways are involved in regulating the functions of GSSs, and thus whether they influence the development of DBM resistance.

## MATERIALS AND METHODS

### DBM strain

Specimens of the *P. xylostella* DBM strain ‘Fuzhou lab’ were reared on radish seedlings without exposure to insecticides for 5 years, spanning at least 100 generations [52].

### Preparation of brush border membrane vesicles

Midgut BBMVs were prepared following the method developed by Wolfersberger et al. [53]. Fifth-instar larvae were immobilized on ice and dissected in cold dissection buffer (17 mM Tris-HCl, pH 7.5, 5 mM ethylene glycol-bis(β-aminoethyl ether)-N,N,N’,N’-tetraacetic acid (EGTA), 300 mM mannitol, 1 mM phenylmethane sulfonyl fluoride (PMSF)) to isolate the midgut epithelium. Midgut epithelial tissue was homogenized in an equal volume of ice-cold 24 mM MgCl_2_, then incubated on ice for 15 min, followed by centrifugation at 25,006 *g* at 4°C for 15 min to collect the supernatant. The centrifuged pellet was resuspended in ice-cold dissection buffer in 0.5 volume of the initial homogenate and then the BBMV extraction procedure was repeated as described above. The supernatants collected from the two extractions were combined and BBMVs were precipitated by centrifugation at 30,000 *g* at 4°C for 1 h and stored at ‒80°C. Protein concentration was measured using the BCA Protein Assay Kit (Rockford, USA) according to the manufacturer’s instructions.

### Modified far-immunoblotting

Ten micrograms of each Bt protein was separated by 10% SDS-PAGE, and transferred onto nitrocellulose membrane using an Amersham Semi-Dry Transfer Unit (Freiburg, Germany). The membrane was stained with Ponceau S, then the band containing the Bt protein was excised and destained. Proteins in the nitrocellulose membrane were then denatured and renatured by gradually reducing the guanidine-HCl concentration [31]. Briefly, proteins were denatured by incubating the membrane in denaturing and renaturing buffer (100 mM NaCl, 20 mM Tris (pH 7.6), 0.5 mM ethylenediaminetetraacetic acid (EDTA), 10% glycerol, 0.1% Tween-20, and 1 mM dithiothreitol (DTT)) containing 6 M guanidine–HCl for 30 min at room temperature. The membrane was then washed with denaturing and renaturing buffer containing 3 M guanidine-HCl for 30 min at room temperature, then washed with denaturing and renaturing buffer containing 0.1 M and no guanidine–HCl for 1 h at 4°C. The membrane was blocked with Pierce protein-free buffer (Rockford, USA) for 1 h at room temperature. Membranes were incubated with 30 μg total BBMV proteins (final concentration 10 μg/mL) in protein-binding buffer (100 mM NaCl, 20 mM Tris (pH 7.6), 0.5 mM EDTA, 10% glycerol, 0.1% Tween-20, and 1 mM DTT) overnight at 4°C. Bt protein on the membrane captures BBMV proteins if they form a complex; the nitrocellulose band containing Bt protein and BBMV proteins was then ready for trypsin digestion.

Far-immunoblot analysis was performed as previously described. Briefly, Bt proteins (Cry1Ac, Cry1Ab, and Cry1Bd) were separated on a 12% SDS-PAGE gel, then transferred to polyvinylidene difluoride (PVDF) membrane. Protein denaturing and renaturing on the membrane was performed exactly as per the protein co-blotting procedure described above. The membrane was then blocked with 5% milk in phosphate-buffered saline (PBS) for 1 h at room temperature. The membrane was then incubated with 5 μg purified His-GSS1 or His-GSS2 proteins (final concentration 1 μg/mL) in protein-binding buffer overnight at 4°C. Membranes were probed with anti-His primary antibodies, then washed with PBS with Tween-20 (PBST), incubated with horseradish peroxidase (HRP)-conjugated secondary antibodies, and exposed to X-ray films after reacting with electrochemiluminescence (ECL) substrates.

### On-membrane digestion

On-membrane digestion was carried out as described by Luque-Garcia et al. [32]. Nitrocellulose bands were washed at least six times with Milli-Q water (Merck Millipore, Shanghai, China), then incubated in trypsin solution (12.5 ng/μl prepared in 50 mM NH_4_HCO_3_ buffer (pH 8)) at 37°C overnight. After digestion, samples were dried in a vacuum, dissolved in acetone (90 μl acetone/4 mm^2^ nitrocellulose), vortexed, and incubated for 30 min at room temperature. Acetone containing dissolved nitrocellulose was carefully removed and precipitated peptides were air-dried. Peptides were resuspended by adding 20 μl of 2% acetonitrile in 0.1% formic acid. All solutions were sonicated for 10 min before mass spectrometry analysis.

### Q Exactive LC MS/MS analysis

In the analysis of complex mixtures, peptides of similar mass often co-elute; therefore, resolution is key in mass spectrometry [54]. Shotgun proteomics using the Q-Exactive instruments (Thermo Fisher Scientific, USA) is usually performed at 17,500 resolution at m/z 200 with a transient length of 60 ms. The higher resolution in MS/MS spectra helps to assign fragments of large precursors. Data were analyzed in MaxQuant using the integrated Mascot search engine [54]. The total number of MS scans exceeded 5000, and the total number of MS/MS scans exceeded 16 000. The average number of isotope patterns detected was close to 35,000, a very high number considering that the gradient was not particularly long, presumably because of the short MS and MS/MS cycle time of 1 s.

### Cloning and purification of GSS1and GSS2

Primers used for cloning in this study are listed in Supplemental Table 3. *GSS1* and *GSS2* were cloned into the pET28A vector using BamHI and HindIII sites, and over-expressed in the BL21 (DE3) *Escherichia coli* strain.

Bacterial cells were harvested by centrifuging at 3000 *g* for 15 min at 4°C. Cells were washed with bacterial cell lysis buffer to remove residual culture medium. Washed cells were harvested by repeating centrifugation at 3000 *g* for 15 min at 4°C. After decanting the supernatant, the wet pellet was weighed and *E. coli* cells resuspended in about 3 mL of lysis buffer per gram of cell pellet. The suspension was stirred for 30 min at 4°C, lysozyme was added to a concentration of 0.1% (*w/v*), then the mixture was incubated for 30 min at 4°C while shaking gently. The suspension was centrifuged at 23,000 *g* for 30 min at 4°C and the supernatant discarded. The pellet was dissolved in inclusion body binding buffer (20 mM Tris-HCL (pH 7.9), 5 mM imidazole, 0.5 M NaCl, 8 M urea). The cleared inclusion bodies solution was applied to a column containing Ni-Agarose beads equilibrated with 3 × 5 mL ultrapure water and 3 × 5 mL protein binding buffer. The column was washed with 15 bed volumes of inclusion body binding buffer before eluting polyhistidine-tagged proteins with 5–10 bed volumes of inclusion body elution buffer (20 mM Tris-HCL (pH 7.9), 500 mM imidazole, 0.5 M NaCl, 8 M urea). Protein purity was typically >90% as determined by SDS-PAGE and Coomassie blue staining. Protein concentrations were measured using the BCA Protein Assay Kit according to the manufacturer’s instructions.

### His-tag pull-down

The Pierce His Tag Protein Interaction Pull-Down Kit (catalog number 21277) was used to detect the binding of GSS1 and GSS2 with Bt proteins. Solubilization of proteins (His-GSS1 and His-GSS2) from inclusion bodies (a requirement of this kit) was carried out according to the method developed by Simpson [55]. Cells were lysed as described in “Cloning and purification of GSS1and GSS2”. The cell lysate was centrifuged at 23,000 *g* for 30 min at 4°C before decanting the supernatant and measuring the wet mass of the pellet. The pellet was resuspended in 10 volumes of lysate washing buffer and the suspension stirred for 1 h at room temperature. The mixture was again centrifuged at 23,000 *g* for 30 min at 4°C, the supernatant decanted and the pellet recovered. Wash steps were repeated three more times. The pellet was then dissolved in 9 volumes of solubilization buffer C per gram wet weight of inclusion body pellet, and the mixture incubated for 1 h at room temperature. Nine volumes of renaturation buffer C was added slowly to the solubilized pellet and the mixture incubated for 2–4 h at 25°C. Two milliliters of Ni-Agarose beads were added to renaturation buffer C and incubated overnight on a magnetic stirrer at 4°C. Protein–bead complexes were collected by pressing renaturation buffer C through a column with a 0.25 nm filter. Polyhistidine-tagged proteins were eluted with 5–10 bed volumes of elution buffer (20 mM Tris-HCL (pH 7.9), 500 mM imidazole, 0.5 M NaCl). Imidazole was removed from purified His-GSS1, His-GSS2 and Bt proteins (Cry1Ac, Cry1Ab and Cry1Bd) by dialysis against Tris-buffered saline buffer (25 mM Tris-HCl, 0.15 M NaCl (pH 7.2)).

A His-tag protein was added to spin columns containing equilibrated HisPur Cobalt Resin (Thermo Scientific, Rockford, USA) and incubated at 4°C for at least 30 min on a rotating platform with a gentle rocking motion. Spin columns were centrifuged at 1250 *g* for 30 s to 1 min to remove solution. Beads were washed 5 times with 400 μL of wash solution. Up to 150 μg of prepared Bt proteins was added to columns, which were then incubated overnight at 4°C. Beads were washed 5 times with washing buffer to remove non-specifically bound proteins. Proteins were eluted by adding 250 μL of elution buffer (1 mL of 290 mM imidazole elution buffer made with 70 μL of 4 M imidazole stock solution to 930 μL of wash solution) to the spin column. Spin columns were incubated for 5 min on a rotating platform with gentle rocking motion, before centrifuging at 1250 *g* for 30 s to 1 min. Proteins were analyzed by SDS-PAGE and visualized by silver stain.

### Plasmids and plant transformation

Plasmids for double-stranded RNA (dsRNA) expression were constructed as previously described [56]. The pBSK intron vector was a pBluescript II SK vector (Stratagene) containing a 120-nucleotide intron of the *Arabidopsis thaliana RTM1* gene between the NotI and XbaI sites. Sense and antisense target fragments with restriction enzyme sites at both ends were obtained by PCR amplifying DBM cDNA clones with primer pairs (Supplemental Table 3). The two PCR fragments were inserted at inverted repeats into the corresponding sites of the pBSK intron vector. The dsRNA construct generated was then used to replace GUS in pBI121 to generate the *Pro*35S::dsRNA construct. The final RNAi vector was introduced into *Agrobacterium tumefaciens* strain GV3101. Transgenic Arabidopsis plants were generated using the floral dip method, screened on half-strength Murashige and Skoog (MS) agar medium containing 30 μg/mL kanamycin.

**Supplemental Table 3.** List of primers

dsRNA expression levels were analyzed using T2 homozygous plants. Freshly hatched DBM larvae were fed with leaves of the *dsGFP* and *dsGSS1* lines for 7 d, respectively. Expressions of *GSS1* and *GSS2* were detected by real-time PCR (primers listed in Supplemental Table 3).

### Quantitative reverse transcription PCR (RT-qPCR)

RT-qPCR was performed using an Eppendorf Mastercycler ep realplex, using gene-specific and allele-specific primers to detect expression patterns. Each reaction was performed in a 20 μL volume containing 10 μL SYBR Green (Fermentas), 0.4 μL Rox Reference Dye II, 1 μL of each primer (10 mM), 1 μL of sample cDNA, and 7.6 μL UltraPure distilled water (Invitrogen). The PCR program used was: 95°C for 10 s, 40 cycles at 95°C for 20 s, 60°C for 30 s. Relative quantification was calculated using the comparative 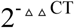 method [57]. All data were normalized to the level of *RP49* from the same sample.

### Plasmids and silkworm transformation

Transformation plasmids were constructed based on the initial piggyBac vectors pBac[3xp3-DsRed, IE1-EGFP]. The 3xp3 promoter was removed by cutting with NotI and AgeI, and was replaced with the HR5 enhancer followed by the IE1 promoter to generate the pBac[HR5IE1-DsRed, IE1-EGFP] plasmid based on homologous recombination using the ClonExpressTM II One Step Cloning Kit (Vazyme Biotech Co., Ltd.). The open reading frames (ORFs) of *GSS1* and *GSS2* were replaced with *EGFP* to generate the silkworm transformation plasmids, pBac[IE1-DsRed, IE1-GSS1] and pBac[IE1-DsRed, IE1-GSS2]. Primers are listed in Supplemental Table 3.

Silkworm microinjection was performed as described by Tan et al. [58]. The transformation plasmid was microinjected into preblastoderm G0 embryos (Nistari strain), which were then incubated at 25°C in a humidified chamber for 10–12 d until larval hatching. Larvae were reared on fresh mulberry leaves or an artificial diet (Nihonnosanko) under standard conditions. Putative transgenic adult G0 were mated with each other, and G1 progeny were scored for the presence of red fluorescence using fluorescence microscopy (Nikon AZ100). Positive G1 progeny were mated with wild-type moths to generate hybrid silkworms expressing one copy of dsRed and a target gene. Hybrid silkworms with red fluorescence were fed with Cry1Bd-treated leaves as described below.

### Bt toxins treatment

To test the tolerance of RNAi-silenced DBM larvae, two leaves from 4-week-old *A. thaliana* plants with genotypes *dsGFP* and *dsGSS1* were laid on a moistened filter paper in a 15 mm petri dish. Freshly hatched DBM larvae were placed on leaves of each genotype to feed for 7 d. On the eighth day, larvae were transferred to fresh leaves coated with either a diluted suspension of Bt toxin in 4-(2-hydroxyethyl)-1-piperazineethanesulfonic acid (HEPES) buffer (pH 8.0), or HEPES buffer as a control. Mortality was recorded after 24 h and LC50 values were calculated by probit analysis based on the dose determined to be high enough to kill 100% of larvae.

To test the tolerance of transgenic silkworms, four squares of mulberry leaf (4 × 4 cm), coated with 50 μL of a diluted suspension of Cry1Bd in HEPES buffer (pH 8.0) were fed to 10 second instar larvae for 24 h. The dose high enough to kill 100% of susceptible larvae was determined. Probit analysis was carried out using SPSS software (Version 12.0) to determine LC50.

## ACKNOWLEDGMENTS

Thanks to: Dr. Minsheng You, Fujian Agriculture and Forestry University, for kindly providing specimens of the *P. xylostella* DBM strain ‘Fuzhou lab’; Dr. Takashi Yamamoto (DuPont Pioneer) for providing Bt proteins; and Yingbo Mao, Shanghai Institutes for Biological Sciences, who kindly provided the pBSK intron vector.

## REFERENCES

1. Bravo A, Likitvivatanavong S, Gill SS, Soberon M. Bacillus thuringiensis: A story of a successful bioinsecticide. Insect Biochem Molec. 2011; 41: 423-431.

2. Bravo A, Gill SS, Soberon M. Mode of action of Bacillus thuringiensis Cry and Cyt toxins and their potential for insect control. Toxicon. 2007; 49: 423–435.

3. Bravo A, Soberon M. How to cope with insect resistance to Bt toxins? Trends Biotechnol. 2008; 26: 573–579.

4. Soberon M, Pardo-Lopez L, Lopez I, Gomez I, Tabashnik BE, Bravo A. Engineering modified Bt toxins to counter insect resistance. Science. 2007; 318: 1640–1642.

5. Bravo A, Gomez I, Conde J, Munoz-Garay C, Sanchez J, Miranda R, Zhuang M, Gill SS, Soberon M. Oligomerization triggers binding of a Bacillus thuringiensis Cry1Ab pore-forming toxin to aminopeptidase N receptor leading to insertion into membrane microdomains. Bba-Biomembranes. 2004; 1667: 38–46.

6. Jurat-Fuentes JL, Adang MJ. Characterization of a Cry1Ac-receptor alkaline phosphatase in susceptible and resistant Heliothis virescens larvae. Eur J Biochem. 2004; 271: 3127–3135.

7. Zhang XB, Candas M, Griko NB, Taussig R, Bulla LA. A mechanism of cell death involving an adenylyl cyclase/PKA signaling pathway is induced by the Cry1Ab toxin of Bacillus thuringiensis. Proc Natl Acad Sci U S A. 2006; 103: 9897–9902.

8. Ferre J, Van Rie J. Biochemistry and genetics of insect resistance to Bacillus thuringiensis. Annu Rev Entomol. 2002; 47: 501–533.

9. Carriere Y, Fabrick JA, Tabashnik BE. Can pyramids and seed mixtures delay resistance to Bt crops? Trends Biotechnol. 2016; 34: 291–302.

10. Furlong MJ, Wright DJ, Dosdall LM. Diamondback moth ecology and management: problems, progress, and prospects. Annu Rev Entomol. 2013; 58: 517–541.

11. Tabashnik BE. Evolution of resistance to Bacillus thuringiensis. Annu Rev Entomol. 1994; 39: 47–79.

12. Tabashnik BE, Liu YB, Finson N, Masson L, Heckel DG. One gene in diamondback moth confers resistance to four Bacillus thuringiensis toxins. Proc Natl Acad Sci U S A. 1997; 94:1640–1644.

13. Tabashnik BE, Malvar T, Liu YB, Finson N, Borthakur D, Shin BS, Park SH, Masson L, de Maagd RA, Bosch D. Cross-resistance of the diamondback moth indicates ered interactions with domain II of Bacillus thuringiensis toxins. Appl Environ Microbiol. 1996; 62: 2839–2844.

14. Heckel DG, Gahan LJ, Baxter SW, Zhao JZ, Shelton AM, Gould F, Tabashnik BE. The diversity of Bt resistance genes in species of Lepidoptera. J Invertebr Pathol. 2007; 95:192–197.

15. Chen Y, Li M, Islam I, You L, Wang Y, Li Z, Ling L, Zeng B, Xu J, Huang Y, Tan A. Allelic-specific expression in relation to Bombyx mori resistance to Bt toxin. Insect Biochem Mol Biol. 2014; 54: 53–60.

16. Guo Z, Kang S, Chen D, Wu Q, Wang S, Xie W, Zhu X, Baxter SW, Zhou X, Jurat-Fuentes JL, et al. MAPK signaling pathway alters expression of midgut ALP and ABCC genes and causes resistance to Bacillus thuringiensis Cry 1 Ac toxin in diamondback moth. PLOS Genet. 2015; 11: e1005124.

17. Atsumi S, Miyamoto K, Yamamoto K, Narukawa J, Kawai S, Sezutsu H, Kobayashi I, Uchino K, Tamura T, Mita K, et al. Single amino acid mutation in an ATP-binding cassette transporter gene causes resistance to Bt toxin Cry 1 Ab in the silkworm, Bombyx mori. Proc Natl Acad Sci U S A. 2012; 109: E1591–E1598.

18. Gahan LJ, Gould F, Heckel DG. Identification of a gene associated with Bt resistance in Heliothis virescens. Science. 2001; 293: 857–860.

19. Gahan LJ, Pauchet Y, Vogel H, Heckel DG. An ABC transporter mutation is correlated with insect resistance to Bacillus thuringiensis Cry1Ac toxin. PLOS Genet. 2010; 6:e1001248.

20. Guo Z, Kang S, Zhu X, Wu Q, Wang S, Xie W, Zhang Y. The midgut cadherin-like gene is not associated with resistance to Bacillus thuringiensis toxin Cry 1 Ac in Plutella xylostella (L.). J Invertebr Pathol. 2015; 126: 21–30.

21. Crickmore C.. Bacillus thuringiensis resistance in Plutella — too many trees? Curr Opin Insect Sci. 2016; 15: 84–88.

22. Candas M, Loseva O, Oppert B, Kosaraju P, Bulla LA Jr. Insect resistance to Bacillus thuringiensis: alterations in the indianmeal moth larval gut proteome. Mol Cell Proteomics. 2003; 2: 19–28.

23. Krishnamoorthy M, Jurat-Fuentes JL, McNall RJ, Andacht T, Adang MJ. Identification of novel Cry1Ac binding proteins in midgut membranes from Heliothis virescens using proteomic analyses. Insect Biochem Mol Biol. 2007; 37: 189–201.

24. Ratzka A, Vogel H, Kliebenstein DJ, Mitchell-Olds T, Kroymann J. Disarming the mustard oil bomb. Proc Natl Acad Sci U S A. 2002; 99: 11223–11228.

25. Ferre J, Real MD, Van Rie J, Jansens S, Peferoen M. Resistance to the Bacillus thuringiensis bioinsecticide in a field population of Plutella xylostella is due to a change in a midgut membrane receptor. Proc Natl Acad Sci U S A. 1991; 88: 5119-5123.

26. Tabashnik BE, Finson N, Johnson MW, Heckel DG. Cross-resistance to Bacillus thuringiensis Toxin CryIF in the diamondback moth (Plutella xylostella). Appl Environ Microbiol 1994; 60: 4627–4629.

27. Tabashnik BE, Finson N, Johnson MW, Moar WJ. Resistance to toxins from Bacillus thuringiensis subsp. kurstaki causes minimal cross-resistance to B. thuringiensis subsp. aizawai in the diamondback moth (Lepidoptera: Plutellidae). Appl Environ Microbiol. 1993; 59: 1332–1335.

28. Liu YB, Tabashnik BE, Meyer SK, Crickmore N. Cross-resistance and stability of resistance to Bacillus thuringiensis toxin Cry1C in diamondback moth. Appl Environ Microbiol. 2001; 67: 3216–3219.

29. Zhao JZ, Li YX, Collins HL, Cao J, Earle ED, Shelton AM. Different cross-resistance patterns in the diamondback moth (Lepidoptera: Plutellidae) resistant to Bacillus thuringiensis toxin Cry1C. J Econ Entomol. 2001; 94: 1547–1552.

30. Pigott CR, Ellar DJ. Role of receptors in Bacillus thuringiensis crystal toxin activity. Microbiol Mol Biol Rev. 2007; 71: 255–281.

31. Wu Y, Li Q, Chen XZ. Detecting protein-protein interactions by far western blotting. Nat Protoc. 2007; 2: 3278–3284.

32. Luque-Garcia JL, Zhou G, Spellman DS, Sun TT, Neubert TA. Analysis of electroblotted proteins by mass spectrometry: protein identification after Western blotting. Mol Cell Proteomics. 2008; 7: 308–314.

33. Nakanishi K, Yaoi K, Nagino Y, Hara H, Kitami M, Atsumi S, Miura N, Sato R. Aminopeptidase N isoforms from the midgut of Bombyx mori and Plutella xylostella - their classification and the factors that determine their binding specificity to Bacillus thuringiensis Cry1A toxin. FEBS Lett. 2002; 519: 215–220.

34. Chang X, Wu Q, Wang S, Wang R, Yang Z, Chen D, Jiao X, Mao Z, Zhang Y. Determining the involvement of two aminopeptidase Ns in the resistance of Plutella xylostella to the Bt toxin Cry1Ac: cloning and study of in vitro function. J Biochem Mol Toxicol. 2012; 26: 60–70.

35. Tabashnik BE, Finson N, Groeters FR, Moar WJ, Johnson MW, Luo K, Adang MJ. Reversal of resistance to Bacillus thuringiensis in Plutella xylostella. Proc Natl Acad Sci U S A. 1994; 91: 4120–4124.

36. Hernandez-Rodriguez CS, Hernandez-Martinez P, Van Rie J, Escriche B, Ferré J. Shared midgut binding sites for Cry1A.105, Cry1Aa, Cry1Ab, Cry1Ac and Cry1Fa proteins from Bacillus thuringiensis in two important corn pests, Ostrinia nubilalis and Spodoptera frugiperda. Plos One 2013; 8: e68164.

37. Carriere Y, Crickmore N, Tabashnik BE. Optimizing pyramided transgenic Bt crops for sustainable pest management. Nat Biotechnol. 2015; 33: 161–168.

38. Ballester V, Granero F, Tabashnik BE, Malvar T, Ferré J. Integrative model for binding of Bacillus thuringiensis toxins in susceptible and resistant larvae of the diamondback moth (Plutella xylostella). Appl Environ Microbiol. 1999; 65: 1413-1419.

39. Krogh A, Larsson B, von Heijne G, Sonnhammer EL. Predicting transmembrane protein topology with a hidden Markov model: application to complete genomes. J Mol Biol. 2001; 305: 567–580.

40. Eisenhaber B, Bork P, Eisenhaber F. Prediction of potential GPI-modification sites in proprotein sequences. J Mol Biol. 1999; 292: 741–758.

41. Cioffi M, Wolfersberger MG. Isolation of separate apical, lateral and basal plasma membrane from cells of an insect epithelium. A procedure based on tissue organization and ultrastructure. Tissue & cell 1983; 15: 781–803.

42. McNall RJ, Adang MJ. Identification of novel Bacillus thuringiensis Cry1Ac binding proteins in Manduca sexta midgut through proteomic analysis. Insect Biochem Mol Biol. 2003; 33: 999–1010.

43. Baxter SW, Zhao JZ, Shelton AM, Vogel H, Heckel DG. Genetic mapping of Bt-toxin binding proteins in a Cry1A-toxin resistant strain of diamondback moth Plutella xylostella. Insect Biochem Mol Biol. 2008; 38: 125–135.

44. Halkier BA, Gershenzon J. Biology and biochemistry of glucosinolates. Annu Rev Plant Biol. 2006; 57: 303–333.

45. Hernández-Martínez P, Navarro-Cerrillo G, Caccia S, de Maagd RA, Moar WJ, Ferré J, Escriche B, Herrero S. Constitutive activation of the midgut response to Bacillus thuringiensis in Bt-resistant Spodoptera exigua. Plos One. 2010; 5: pii: e12795.

46. McNall RJ. Proteomic analyses of the interaction of insect midgut proteins with Bacillus thuringiensis toxins. 2014; PhD thesis, Univ Georgia.

47. Yamazaki T, Hori H, Ishikawa T, Pandian GN, Okazaki K, Haginoya K, Tachikawa Y, Mitsui T, Miyamoto K, Angusthanasombat C. Midgut juice of Plutella xylostella highly resistant to Bacillus thuringiensis Cry1Ac contains a three times larger amount of glucosinolate sulfatase which binds to Cry1Ac compared to that of susceptible strain. Pestic Biochem Physiol. 2011; 101: 125–131.

48. Hanson SR, Best MD, Wong CH. Sulfatases: structure, mechanism, biological activity, inhibition, and synthetic utility. Angewandte Chemie. 2004; 43: 5736–5763.

49. Derbyshire DJ, Ellar DJ, Li J. Crystallization of the Bacillus thuringiensis toxin Cry1Ac and its complex with the receptor ligand N-acetyl-D-galactosamine. Acta Crystallogr D Biol Crystallogr. 2001; 57: 1938-1944.

50. Dhoot GK, Gustafsson MK, Ai X, Sun W, Standiford DM, Emerson CP Jr. Regulation of Wnt signaling and embryo patterning by an extracellular sulfatase. Science, 2001; 293: 1663–1666.

51. Ragsdale EJ, Muller MR, Rodelsperger C, Sommer RJ. A developmental switch coupled to the evolution of plasticity acts through a sulfatase. Cell, 2013; 155: 922-933.

52. You M, Yue Z, He W, Yang X, Yang G, Xie M, Zhan D, Baxter SW, Vasseur L, Gurr GM, et al. A heterozygous moth genome provides insights into herbivory and detoxification. Nat Genet. 2013; 45: 220–225.

53. Wolfersberger M. Preparation and partial characterization of amino acid transporting brush border membrane vesicles from the larval midgut of the cabbage butterfly. Comparative Biochemistry and Physiology Part A: Physiology 1987; 86: 301–308.

54. Michalski A, Damoc E, Hauschild JP, Lange O, Wieghaus A, Makarov A, Nagaraj N, Cox J, Mann M, Horning S. Mass spectrometry-based proteomics using Q Exactive, a high-performance benchtop quadrupole Orbitrap mass spectrometer. Mol Cell Proteomics. 2011; 10: M111. 011015.

55. Simpson RJ. Solubilization of Escherichia coli recombinant proteins from inclusion bodies. Cold Spring Harb Protoc. 2010; doi:10.1101/pdb.prot5485.

56. Mao YB, Tao XY, Xue, XY, Wang LJ, Chen XY. Cotton plants expressing CYP6AE14 double-stranded RNA show enhanced resistance to bollworms. Transgenic Res. 2011; 20: 665–673.

57. Schmittgen TD, Livak KJ. Analyzing real-time PCR data by the comparative C(T) method. Nat Protoc. 2008; 3: 1101–1108.

58. Tan A, Fu G, Jin L, Guo Q, Li Z, Niu B, Meng Z, Morrison NI, Alphey L, Huang Y. Transgene-based, female-specific lethality system for genetic sexing of the silkworm, Bombyxmori. Proc Natl Acad Sci U S A 2013; 110: 6766–6770.

